# The neglected part of the microbiome: Prophage TJ1 regulates the bacterial community of the metaorganism *Hydra*

**DOI:** 10.1101/607325

**Authors:** Janina Lange, Sebastian Fraune, Thomas C.G. Bosch, Tim Lachnit

**Affiliations:** Zoological Institute, Christian-Albrechts-University Kiel, 24118 Kiel, Germany

**Keywords:** Holobiont, microbiome, host-microbe interaction, virus

## Abstract

Many multicellular organisms are closely associated with a specific bacterial community and therefore considered “metaorganisms”. Controlling the bacterial community composition is essential for the stability and function of metaorganisms, but the factors contributing to the maintenance of host specific bacterial colonization are poorly understood. Here we demonstrate that in *Hydra* the most dominant bacterial colonizer *Curvibacter sp.* is associated with an intact prophage which can be induced by different environmental stressors both *in vitro and in vivo*. Differences in the induction capacity of *Curvibacter* phage TJ1 in culture (*in vitro)* and on *Hydra* (*in vivo*) imply that the habitat of the prokaryotic host and/or bacterial frequency dependent factors influence phage inducibility. Moreover, we show that phage TJ1 features a broad host range against other bacterial colonizer and is directly capable to affect bacterial colonization on *Hydra.* From these results we conclude that prophages are hidden part of the microbiome interfering with bacteria-bacteria interactions and have the potential to influence the composition of host associated bacterial communities.

## Introduction

Eukaryotic organisms are living in a close relationship with a complex microbial community, composed of bacteria, fungi, viruses and protists. This close association can be beneficial for both partners and forms a complex unit termed metaorganism or holobiont (Bosch and McFall-Ngai, 2011; Rosenberg *et al.*, 2007). Disturbance or loss of the natural associated bacterial community can facilitate the invasion of pathogens and lead to reduced host fitness (Van Rensburg *et al.*, 2015; Bates *et al.*, 2018; Fraune *et al.*, 2014).

Host genetics and innate immunity play an important role in establishing and maintaining the microbial composition of metaorganisms. For some animal species including *Hydra* it has been shown that the eukaryotic host actively selects and shapes its specific bacterial community (Franzenburg *et al.*, 2013a; Goodrich *et al.*, 2014; Franzenburg *et al.*, 2012; Pietschke *et al.*, 2017). In *Hydra*, bacteria-bacteria interactions are an important component contributing to the fitness of the metaorganism (Fraune *et al.*, 2014). The most abundant bacterial colonizer of *Hydra vulgaris* (AEP) is the proteobacteria *Curvibacter sp.*. Interestingly, *Curvibacter* can protect the animal host from fungal infection if one of the other main colonizers such as *Duganella sp.* (11%), *Undibacterium* sp. (2.1%) or *Pelomonas* sp. (0.2 %) is also present (Fraune *et al.*, 2014). Co-occurrence of these bacteria is thus essential to provide this beneficial antifungal host defense. Li et al. (2015) investigated the interaction between the two most abundant bacterial colonizers of *Hydra, Curvibacter* sp. and *Duganella* sp., *in vitro* and observed that in mono-culture *Duganella sp.* features higher growth rate than *Curvibacter* sp.. In contrast to the expectation that *Duganella sp.* would outcompete *Curvibacter* sp. in co-culture experiments, a frequency dependent suppression of *Duganella* sp. was detected (Li *et al.*, 2015). The observation of a frequency dependent growth rate indicates that the interactions among bacteria in co-culture are beyond a simple case of direct competition and it has been predicted by modelling that this interaction is mediated by a temperate phage integrated in the genome of *Curvibacte*r sp. (prophage) (Li *et al.*, 2017). The fresh water polyp *Hydra* is not only associated with a host specific bacterial community (Franzenburg *et al.*, 2013b) but also features a host specific viral community of which more than 50% are bacteriophages (Grasis *et al.*, 2014). Due to the fact that phages are often obligate killers to their host cells but also to other bacteria, they have a strong selective effect on bacterial populations and are hypothesized to shape whole bacterial communities (Bohannan and Lenski, 2000; Gómez and Buckling, 2011; Suttle, 2007; Koskella and Meaden, 2013). While lytic phages infect bacteria, get multiplied and kill their bacterial host (Lenski, 1988), temperate phages can undergo lysogenic conversion, where the phage genome is replicated along with the genome of its host and is transferred vertically (De Paepe *et al.*, 2014). By this they can increase the fitness of its bacterial host by altering the bacterial geno- and phenotype (Oakey and Owens, 2000). During stress for the bacterial host cell, temperate phages can switch from lysogenic to lytic cycle (prophage induction) (Ranquet *et al.*, 2005; Nanda *et al.*, 2015). Free phages are then released to the environment and are able to cross-infect and kill other bacteria (Livny and Friedman, 2004; Refardt, 2012). In this manner excised prophages may function as weapons in competition with other (susceptible) bacteria (Bossi *et al.*, 2003; Burns *et al.*, 2015). In a previous study we could demonstrate that the prophage of *Curvibacter* sp. is inducible and can lytically infect the second most abundant colonizer of *Hydra Duganella sp*. *in vitro*. In the present study we further investigate the function of *Curvibacter* phage (TJ1) in the host context and hypothesized that phage TJ1 interferes with bacteria-bacteria interactions and has the potential to influence the composition of host associated bacterial community. To test this hypothesis, we investigated the presence and induction capacity of the prophage associated with *Curvibacter* sp. and its ability to cross-infect and downregulate other *Hydra* associated bacteria in culture (*in vitro*) and on *Hydra* (*in vivo*).

Our results demonstrate that *Curvibacter* phage (TJ1) can be induced in culture (*in vitro)* and on *Hydra (in vivo)*. In association with *Hydra, Curvibacter* sp. was more susceptible for phage induction at reduced water temperature, suggesting an impact of the host environment on phage bacteria interactions. Finally, we could demonstrate that phage TJ1 has a broad host range against other bacterial colonizers and has the potential to regulate bacterial colonization on *Hydra.*

## Material and Methods

### Identification and induction of *Curvibacter* prophage

First, we screened the bacterial genome of *Curvibacter* sp. (strain AEP1.3; GenBank:CP015698.1) for the presence of prophage signatures using the online software PHASTER (Phage Search Tool Enhanced Release) (Arndt *et al.*, 2016). Second, we used Mitomycin C assay to test the induction capacity of the phage. Therefore, *Curvibacter* sp. was grown in 3% (w/v) R2A broth media (Sigma-Aldrich) under shaking conditions at 250 rpm at 18°C. Exponentially growing bacterial overnight cultures were inoculated with 0.05 µg/ml Mitomycin C to induce phage replication (Sekiguchi and Takagi, 1960). After an incubation time of 16 h bacterial cells were removed by two consecutive low-speed-centrifugation steps at 4,266 × g in a ThermoScientific Heraeus Multifuge 3SR at 4°C for 30 min. The supernatant was filtered 0.2 µm and phage particles were pelleted by ultracentrifugation at 25,000 rpm (72,700 × g) in a Beckman 45 Ti rotor at 4°C for 2h. The pellet was re-suspended in 3 ml SM-Buffer (50mM Tris; 100 mM NaCl; 8 mM MgSO_4_; pH 7.5). Re-suspended phages were layered onto a pre-formed Cesium chloride gradient consisting of the densities (2ml: 1.7; 1.5; 1.3; 1.2 g/cm^3^ and 1 ml 1.1 g/cm^3^) in SM-buffer and centrifuged at 28.000 rpm (135,000 × g) in a Beckman SW 41 rotor at 4°C for 2 h. The band containing phages was removed by syringe, diluted 1:3 in SM-Buffer and pelleted by centrifugation at 22 000 rpm (83,000 × g) in a Beckman SW 41 rotor at 4°C for 2 h. Phages were re-suspended in 200 µl SM-Buffer and stored at –80°C until DNA extraction. Sub-samples of phages (5 µl) were further characterized morphologically by negative staining in 2% (w/v) aqueous uranyl acetate and visualized by transmission electron microscopy (TEM) using a Technai Bio TWIN at 80 kV and a magnification of 40 000–100 000.

### Phage DNA extraction and Sequencing

Phage DNA was extracted from two-hundred microliters of purified phages according to the protocol developed by Thurber and colleagues (Thurber *et al.*, 2009) with minor modifications. In brief, 22µl 2M Tris-HCL (pH 8.5)/0.2 M EDTA, 10 µl 0.5 M EDTA and 268 µl formamid were added to the samples and incubated for 30 min at room temperature. DNA was precipitated by adding two volumes of ethanol and incubation at −20°C overnight. DNA was pelleted by centrifugation at 13,000 × g at 4°C for 20 min. The pellet was washed with 70% (v/v) ethanol. 100 µl SDS extraction buffer (1% SDS; 100 mM Tris; 20 mM EDTA; pH 7.5), 1% 2-mercaptoethanol and proteinase K (0.5 mg/ml) were added to the pellet and incubated for 30 min at 36°C and 15 min at 56°C. Twice the amount of DNA-extraction buffer (100 mM Tris, pH 8.0; 1.4 M NaCl; 20 mM EDTA; 1% 2-mercaptoethanol; 2% (w:v) CTAB) were added followed by an incubation step at 65°C for 10 min. An equal volume of chloroform:isoamyl alcohol (24:1) was added to the warm solution, mixed thoroughly and centrifuged at 13,000 × *g* at room temperature for 5 min. Supernatant was transferred into a new tube and DNA was precipitated by the addition of 0.7 volume of isopropanol. After an incubation at −20°C for 2 h DNA was pelleted by centrifugation at 13,000 × g at 4°C for 20 min. DNA was washed with 70% (v/v) ethanol and dissolved in 50 µl water. Nextera XT kit (Illumina) was used for library preparation and 2 × 150 bp paired-end sequencing was conducted on a MiSeq platform (Illumina) at Centre for molecular biology in Kiel. Trimmomatic V.0.36 (Bolger *et al.*, 2014) was used for sequence adaptor removal and read trimming. Trimmed and quality controlled reads were finally assembled using SPAdes V.3.1.11 (Bankevich *et al.*, 2012). The assembled *Curvibacter* phage TJ1 genome is publically available under the GenBank accession number MH766655 in the NCBI database.

### Phage annotation and comparison

Phage TJ1 was annotated using the genome annotation service RAST (Rapid Annotation using Subsystem Technology). The phage capsid protein was compared with different phages by Psi-Blast. The most closely related was *Burkholderia* phage KS10 (NCBI: Reference Sequence: NC_011216.1) by SEED Viewer (Wattam *et al.*, 2017). Gene organization and sequence similarity between *Curvibacter* phage TJ1 and *Burkholderia* phage KS10 was graphically illustrated by SEED Viewer centered on the phage capsid protein (Wattam *et al.*, 2017).

In a next step we controlled RAST predicted ORFs by GeneMark (Borodovsky and McIninch, 1993) and conducted similarity searches by BLASTP (Altschul *et al.*, 1997), SWISS-PROT (Bairoch and Apweiler, 2000), UniProt (2019) and Pfam databases (El-Gebali *et al.*, 2019). Based on BLASTP results we selectively downloaded bacterial genomes from the National Centre for Biotechnology Information (NCBI) and screened them for the presence of prophages by PHASTER (Phage Search Tool Enhanced Release) (Arndt *et al.*, 2016). Prophage sequences were extracted and submitted together with the sequence of Burkholderia phage KS10 to VICTOR (Meier-Kolthoff and Göker, 2017) for phylogenetic analysis using the GENOME-BLAST Distance Phylogeny method (GBDP) (Meier-Kolthoff and Göker, 2017).

### Phage quantification

Phages were quantified by qPCR (Imamovic *et al.*, 2010; Refardt, 2012). In order to take into account that phage sequences derive from either induced phages or from bacterial cells carrying integrated prophages in their genome we quantified both phage TJ1 and bacteria via qPCR. We calculated the ratio between phage and bacteria to exclude the effect of differences in growth or polyp size. For the quantification of phage TJ1 we designed a specific set of primers targeting the phage tail gene (F: 5’-GCTTTGACCTGTCGTTCATCC-3’ and R: 5’-CGGGTTTGTTGGATAGGTCGT-3’). For bacteria quantification we used either primer specific for the recA gene of *Curvibacter* sp. (strain AEP1.3) (F: 5’-TTCGGCAAGGGCACCATC-3’ and R: 5’-ACGACTCCGGGCCATAGA-3’) or Eubacterium Primer (F: 5’-CCTACGGGAGGCAGCAG-3’ and R: 5’-ATTACCGCGGCTGCTGGC-3’) for quantification of the other bacterial colonizer (cross-infection experiment *in vivo*). Tests of primer sets on DNA extracts of germ free control polyps were negative. QPCR reactions were performed in a 25 μl volume with 12.5 μl GoTaq qPCR MasterMix (Promega), 10 pmol/µl forward and reverse primers and 10 ng/µl template DNA. Cycler (Applied Biosystems 7300 Real Time PCR System) conditions were as follows: 95 °C for 2 min, 40 cycles of 95 °C for 15 s, 59 °C for 30 s, 60 °C for 35 s, followed by a dissociation step. At the end of each cycle at 60°C the fluorescence was measured. Sequence data are deposited at Sequence Read Archive (BioSample accessions: SAMN11334657-SAMN11334682).

### Environmental stress induction of phage TJ1 *in vitro*

Bacterial log phase cultures were grown in R2A media until they reached an optical density (OD600) of 0.26. Subsequently 6 ml bacterial cultures were exposed separately and with replication (n = 5) to one of the following conditions, while the rest of the environmental conditions remain normal: altered temperature (23°C, 12°C), elevated pH (9.5, 8.5), higher nutrition (4 × R2A medium (12 g/l)) or normal conditions (18°C, pH 7) serving as control. After an incubation time of 16 h, DNA was extracted using the DNA blood and tissue kit (Qiagen) followed by phage and *Curvibacter* sp. quantification by qPCR (see above).

### Recolonization of *Hydra*

Experiments were carried out using *Hydra vulgaris* (AEP) (Hemmrich *et al.*, 2007). Prior to recolonization animals were cultured under constant laboratory conditions including *Hydra* culture medium (0.28 mM CaCl_2_, 0.33 mM MgSO_4_, 0.5 mM NaHCO_3_ and 0.08 mM KCO_3_), food (first instar larvae of *Artemia salina*, fed four times per week) and constant water temperature at 18°C, according to standard procedures (Bosch *et al.*, 1988). Germfree *Hydra* were generated by exposing animals to an antibiotic cocktail containing 50 µg/ml of Ampicillin, Rifampicin, Streptomycin, Spectinomycin and Neomycin as previously described (Franzenburg *et al.*, 2012). Antibiotic solutions were exchanged every second day. After 2 weeks of antibiotic treatment animals were transferred to antibiotic free sterile *Hydra* culture medium for 3 days. Sterility was controlled with previous established methods (Franzenburg *et al.*, 2013b). 5000 CFU/ml of the corresponding bacterial strain was added to the surrounding water (50 ml) of germfree polyps and incubated for one day. Afterwards polyps were washed in sterile *Hydra* medium in order to remove unattached bacteria from the surrounding medium and transferred into new container with fresh, sterile *Hydra* medium and used for subsequent experiments. Depending on the bacterial strain initial abundance of monocolonised *Hydra* was roughly 15-400 CFU per polyp.

### Environmental stress induction of phage TJ1 *in vivo*

*Curvibacter* sp. monocolonized *Hydra* polyps (see above) were transferred to 1.5 ml tubes with 200 µl *Hydra* culture medium. After 3 days of settlement they were exposed separately and with replication (n = 5) to one of the following conditions, while the rest of the environmental conditions remain normal: altered temperature (23°C, 12°C), higher pH (9.5, 8.5) and elevated nutrition (10% R2A diluted in sterile *Hydra* culture medium) or normal conditions (18°C, pH 7) serving as control. After 16 h incubation time DNA was extracted from the polyp including surrounding water. Phages and *Curvibacter* were quantified by qPCR.

### Cross-infection and host range assay *in vitro*

Cross-infectivity of phage TJ1 was tested by spot assays in double agar layer (Adams, 1959) against our bacterial culture collection consisting of diverse bacterial strains isolated from different *Hydra* species (see Table 1). Exponentially growing bacterial strains were mixed into 5 ml preheated top-agar (R2A medium, 5 mM CaCl_2_, 5 mM MgCl_2_, 0.4% agarose) and poured onto R2A agar plates (R2A medium, 1.5% agar). Plates were dried for at least 30 min and 10 µl of purified phages and 10 µl sterile *Hydra* culture medium as control were spotted onto each bacterial lawn (n = 3). Plaque formation was checked after incubation at 18°C for 12-24 h. Purity of phages was controlled for the presence of potential bacterial contaminations by plating out the phage suspension on R2A agar plates.

### Cross-infection assay *in vivo*

This experiment was conducted with a subset of bacterial strains specific for the colonization of *Hydra vulgaris* (AEP). Germfree *Hydra vulgaris* (AEP) polyps were monocolonized with *Curvibacter* sp., *Duganella* sp., *Undibacterium* sp., *Acidovorax* sp. *or Pelomonas* sp. (see above) in log-phase.After 3 days of settlement, the recolonized polyps were transferred separately to 1.5 ml tubes, containing 200 µl sterile *Hydra* culture medium. Phage TJ1 was purified from 100 ml culture and resuspended in sterile *Hydra* culture medium andwere then added to 5 monocolonized polyps at a concentration of 1 ×10^4^ PFU/ml. As control served 5 monocolonized polyps that received an equal volume of pure sterile *Hydra* culture medium. After 0 h, 24 h and 72 h post infection the amount of colonizing bacteria and the amount of phages were quantified. For bacterial quantification polyps were homogenized separately, diluted and plated out onto R2A agar plates (n = 5) and incubated at 18°C. CFU per polyp were counted after 2-4 days of incubation. For the quantification of phages that were released to the surrounding water, DNA was extracted from the polyps and their surrounding sterile *Hydra* culture medium, using the DNA Blood and Tissue Kit (Qiagen). Phages and bacteria were quantified by qPCR (see above).

### Phage induction by different bacterial colonizers *in vivo*

In order to test if phage TJ1 is already active or can be induced only in the presence of different bacterial colonizers, *Hydra* polyps were monocolonized with exponentially growing *Curvibacter* and after three days of settlement they were transferred to 1.5 ml tubes. The polyps were subsequently exposed separately and with replication (n = 5) to 5000 CFU/ml to one of the five main colonizers: *Curvibacter* sp., *Duganella* sp., *Undibacterium* sp., *Pelomonas* sp. and *Acidovorax* sp. After 16 h incubation time a DNA extraction of the polyp and its surrounding water was conducted and the bacteria and phages were quantified via qPCR (see above).

### Impact of phage TJ1 on microbial community

Germfree *Hydra vulgaris* (AEP) polyps were colonized with *Duganella* sp., *Undibacterium* sp., *Acidovorax* sp. and *Pelomonas* sp. (see above). After three days of settlement we added phage TJ1 at a concentration of 1×10^4^ PFU/ml to the recolonized polyps and sterile S-Medium to polyps that served as control with replication (n=5). After 24 h and 72 h we extracted DNA described above. Bacterial community composition was analyzed by amplicon sequencing of the variable region V1-V2 of the 16S rRNA gene using the forward primer 27F (5′-AATGATACGGCGACCACCGAGATCTACAC XXXXXXXX TATGGTAATTGT AGAGTTTGATCCTGGCTCAG-3′) and reverse primer 338R (5′-CAAGCAGAAGACGGCATACGAGAT XXXXXXXX AGTCAGTCAGCC TGCTGCCTCCCGTAGGAGT-3′) containing the Illumina adaptor p5 (forward) and p7 (reverse) and unique MIDs (designated as XXXXXXXX). PCR reactions were performed in duplicates using Phusion Hot Start DNA Polymerase (Finnzymes, Esppoo, Finnland). PCR cycling conditions were: 98°C for 30 s, 30 × [98°C – 9 s, 55°C – 30 s, 72°C – 90 s], 72°C – 10 min. PCR products were combined and purified by MinELute Gel Extraction Kit (Qiagen) after agarose gel electrophoresis. Sequencing was performed on the Illumina MiSeq platform at the sequencing facility of the Kiel Institute for Clinical Molecular Biology (IKMB). Sequence data were analysed using the MOTHUR packages(Schloss *et al.*, 2009) according to the MISeq SOP (Kozich *et al.*, 2013). In Brief, MiSeq paired-end reads were assembled and quality controlled finally resulting in 9837 sequences per sample. Sequences were grouped into operational taxonomic units (OTU) using a 97 % similarity threshold. Sequences were aligned to SILVA 128 Database and taxonomically classified by RDP classifier.

### Statistical data analyzes

All statistical analyzes were conducted using the statistic program R® (version 3.2.3 (2015-12-10) “Wooden Christmas-Tree” (Copyright (C) 2015 The R Foundation for Statistical)). The Levene’s test was used to check for homogeneity of variance. Some data did not fulfill the criteria of normality. Therefore, the effect of environmental stressors on phage induction was analyzed by using a Wilcoxon-Rank-Test. The amount of phages over time and the effect of other colonizers on prophage induction were analyzed by using a generalized linear model (GLM), with a following Tukey HSD post-hoc test for multiple comparisons or pairwise comparisons. The statistical analysis of the cross-infection *in vivo* was conducted by ANOVA, with a following Tukey HSD post-hoc test for multiple comparisons.

## Results and Discussion

### *Curvibacter* sp. (strain AEP1.3) harbours a prophage that can be induced in culture and on *Hydra*

Bio-computational analysis of the genome of *Curvibacter* sp. (Pietschke *et al.*, 2017) (strain AEP1.3) predicted an intact prophage within the region 444 644-484 825 bp and an incomplete prophage within the region (547656-566074). To test the induction capacity and to visualize the *Curvibacter* phage TJ1, we first induced the phage with Mitomycin C. Low concentrations of Mitomycin C between 0.05 and 0.1 µg/ml were sufficient to reduce the growth of the bacterial host *Curvibacter* (Fig. 1A). The decline in growth is an indicator for phage induction and replication. We verified the presence of phage TJ1 by gradient ultracentrifugation, negative staining with uranyl acetate and visualization via transmission electron microscopy (TEM) (Fig. 1B). Transmission electron micrographs displayed morphological similarity of phage TJ1 to *Myoviridae* featuring an isometric head of 50 nm in diameter and a contractile tail of 80 by 20 nm with small tail fibers (Fig. 1B).

**Fig 1.**
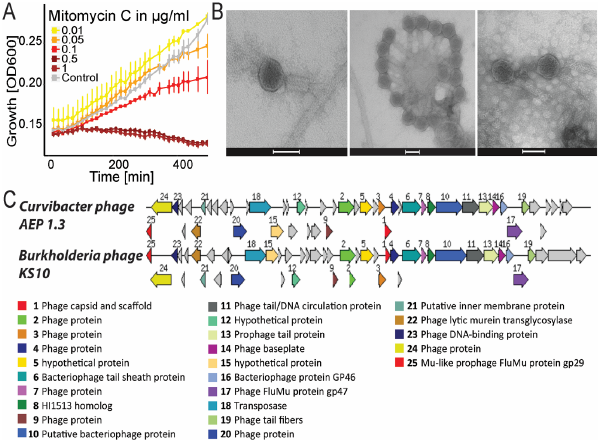
**A:** Mean growth (OD600) (± SE) of *Curvibacter* sp. without treatment (Control, grey) and when exposed to 5 different concentrations of Mitomycin C (n=3)**. B:** Transmission Electron micrographs of *Curvibacter* phage TJ1 negative stained with 2% (w/v) aqueous uranyl acetate. Scale bars represent 50 nm. **C:** Annotation overview of phage TJ1 compared to *Burkholderia* phage KS10 (NCBI: Reference Sequence: NC_011216.1). The graphic is centered on the phage capsid coding region colored in red and numbered with 1. Sets of genes with similar sequence are grouped with the same number and color.

Sequencing the linear double-stranded DNA of phage TJ1 (Supplementary Figure S1) revealed that the 37 079 bp long phage genome featured 54 proteins including capsid protein, phage tail, tail sheath and tail fiber proteins as well as transposase and lytic murein transglycosylase (Table S1). The second prophage seems to be inactive as we did not observe different morphotypes by TEM or detected it in our sequencing data. A comparison of phage TJ1 to other known phages indicated highest similarity to *Burkholderia* phage KS10 with 25 of proteins in common (Fig. 1C). Similarity searches of phage TJ1 by BLASTP revealed the presence of TJ1 predicted proteins in several other bacterial strains including Betaproteobacteria but also Gammaproteobacteria. Screening the genomes of these bacteria by PHASTER we could reveal that these proteins were located within prophages. Phylogenetic analysis of phage sequences by VICTOR using the GENOME-BLAST Distance Phylogenetic method (GBDP) demonstrated the presence of phylogenetic similar phages in distantly related bacteria implicating a brought host range of these phages infecting Betaproteobacteria as well as Gammaproteobacteria (Supplementary Fig. S2).

To test if the prophage of *Curvibacter* sp. can be excised from the host genome under more natural conditions we tested the impact of different environmental factors including temperature, pH and nutrition on prophage induction both in culture (*in vitro*) (Fig. 2A) and in association with *Hydra* (*in vivo*) (Fig. 2B). We quantified phages and *Curvibacter* via qPCR (Fig. S3, Fig. S4) and calculated phage/*Curvibacter* ratio to exclude the effect of different colonization successes which highly influenced by the polyp size. When *Curvibacter* sp. was grown in bacterial growth media in the absence of the eukaryotic host, environmental stressors, like altered temperature, higher pH or higher nutrition had no significant effect on prophage induction in comparison to the control. Phage TJ1 could only be induced by Mitomycin C (Wilcoxon-Rank-Sum-test: W=0, p=0.007) (Fig. 2C). Living in association with *Hydra* phage TJ1 also be induced by Mitomycin C (Wilcoxon-Rank-Sum-test: W=0, p=0.01193 (Fig. 2D). The amount of phage TJ1 produced under Mitomycin C induction was significantly higher when its bacterial host *Curvibacter* was living in association with the *Hydra* epithelium compared to the *in vitro* studies (One-Way-ANOVA, F=10.13 p=0.002). Interestingly a temperature reduction from 18°C to 12°C affected the number of induced phages only if the bacterial host *Curvibacter sp.* lived in associated with *Hydra* (Wilcoxon-Rank-Sum-test: W=25, p =0.01193). Changes in pH-values, elevated temperature or nutrition did not lead to a significant prophage induction, neither *in vitro*, nor *in vivo* (Fig. 2B, 2D). This observation suggests that lysogenesis is maintained and phages are not induced under a natural range of variable environmental condition. As phage TJ1 was not detectable in the virome data set of *Hydra* (Grasis *et al.*, 2014), ambient or elevated water temperature also seems to have no effect on prophage induction on polyps, that are associated with a complex, natural bacterial community.

**Fig 2.**
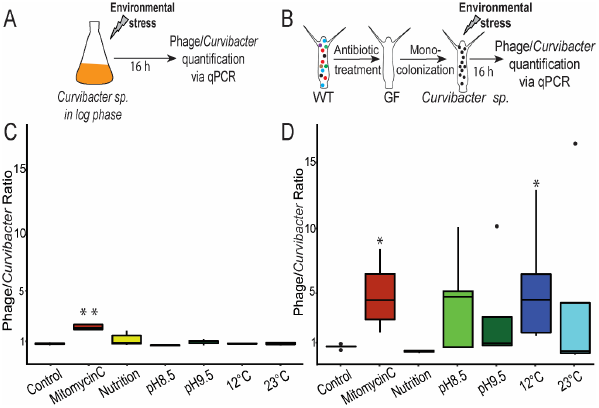
**A:** *In vitro* experimental set-up. *Curvibacter* sp. were grown at log phase and exposed to several environmental stressors. Prophage induction was quantified via qPCR (n=5). **B:** *In vivo* experimental set-up. WT polyps were treated with antibiotics to generate germfree *Hydra.* These germfree polyps were recolonized in mono-association with *Curvibacter* and exposed to different environmental stressors. Prophage induction was quantified via qPCR (n=5); **C:** Boxplot of phage/*Curvibacter* ratio after exposure to the environmental stressors. Stars (*=p<0.05, **= p<0.01) indicate significantly differences between treatment and control (Wilcoxon rank sum test). **D:** Boxplot of phage/*Curvibacter* ratios after the polyps were exposed to the environmental stressors. Stars (*=p<0.05, **= p<0.01) indicate significantly differences between treatment and control (Wilcoxon rank sum test).

The observation that the prophage can be induced *in vivo* by lowering the water temperature is interesting since temperature serves as an important environmental cue in *Hydra* and can affect many developmental processes including body size and budding (Bode *et al.*, 1973; Bisbee, 1973; Stiven, 1965) as well as onset of sexual reproduction (Littlefield *et al.*, 1991).

Prophage excision is expected to be strongest when bacteria are exponentially growing (Madera *et al.*, 2009), due to the high expression of genes involved in the SOS response of the bacterium (Nanda *et al.*, 2014). In the glycocalyx of *Hydra* epithelia, we would expect decreased bacterial growth compared to liquid culture conditions due to nutrient limitation and thereof a reduction of phage inducibility. However, we observed the opposite; phage inducibility was significantly higher (One-Way-ANOVA, F=10.13 p=0.002) when bacteria were associated with their host. The observed differences in the induction capacity add support to the view that the host environment has a strong influence on phage bacteria interactions (De Paepe *et al.*, 2016). Important factors in the epithelial-derived mucus layer may include nutrients as well as components of the innate immune system (Deines *et al.*, 2017; Pietschke *et al.*, 2017). However, there are also other factors that might play a role in the differential induction capacity. Firstly, we cannot exclude a possible additive effect of residual antibiotics on prophage induction which were used in the process of generating germfree polyps, although we stopped antibiotic treatment 6 days before the experiments and washed polyps twice in sterile *Hydra* medium before they were used for all experiments. Secondly, dilution or density dependent issues may lead to a lower induction rate *in vitro* compared to *in vivo* conditions. Recently we predicted by mathematical modelling that at low frequencies the main rout of phage TJ1 production is via the lytic pathway, while at high densities the lysogenic cycle is favored (Li *et al.*, 2017).

### *Curvibacter* phage TJ1 can infect a broad range of *Hydra* associated bacteria

To test the ability of phage TJ1 to cross-infect other bacteria we spotted purified phages onto a lawn of *Hydra* associated bacteria and used plaque formation as sign of infection success. For a variety of bacteria belonging to Betaproteobacteria we could observe plaque formation, while no plaque formation was visible for Gammaproteobacteria and *Bacteroidetes* (Supplementary Table S2). Phage TJ1 could not only infect closely related bacterial strains affiliating to the same family as their original bacterial host (*Comamonadaceae*) but seems to be lytic in a broad spectrum of bacteria. This includes *Undibacterium* sp. and *Duganella* sp., (*Oxalobacteraceae)*, as well as *Vogesella* sp., a less abundant bacterium of *Hydra*, belonging to the order *Neisseriales* (Supplementary Table S2). To examine whether phage TJ1 can cross-infect bacteria not only *in vitro* but also when they are living on *Hydra*, we monocolonized germfree *Hydra* with the most abundant bacterial colonizer: *Curvibacter* sp., *Duganella* sp., *Undibacterium* sp., *Acidovorax* sp. and *Pelomonas* sp. (Fraune *et al.*, 2014) and subsequently inoculated them with phage TJ1. After 0 h, 24 h and 72 h post phage infection we quantified living bacterial cells by counting (CFU/polyp) and estimated phage/bacteria ratio by qPCR (Fig. 3A). While the phage did not interfere with the growth of *Curvibacter* sp. and *Acidovorax* sp., we observed a significant reduction of *Duganella* sp. and *Undibacterium* sp. After 72 h post phage exposure (One-way ANOVA, *Duganella:* F= 4.111; p<0.006, *Undibacterium*: F=3.973; p=0.013) (Fig. 3B). Interestingly we observed a significant reduction of CFUs in polyps that were monocolonized with *Pelomonas* sp. 72 h post phage infection (One-way ANOVA, F = 22.92, p< 0.001) (Fig. 3B), although *Pelomonas* sp. could not be infected by phage TJ1 in *in vitro* experiments (Supplementary Table 1). The impact of phage TJ1 on bacterial numbers (Fig. 3B) was accompanied by an elevated amount of phages (Fig. 3C). The number of phages increased significantly over time in polyps which were recolonized with *Undibacterium* (GLM= 2.7373, p= 0.0103), *Duganella* (GLM, F= 2.7373, p=0.0246) and *Pelomonas* (GLM, F = 2.7373, 0-24h: p= 0.0011; 0-72h: p< 0.0024)) (Fig. 3C). This implies a successful infection and replication of phage TJ1. There was no significant increase of phages detectable in polyps recolonized with *Curvibacter* sp. or *Acidovorax* sp. (Fig. 3C).

**Fig 3.**
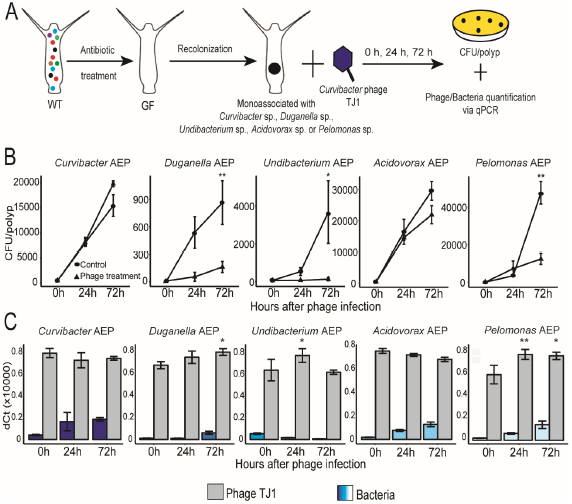
**A:** Experimental set-up. WT polyps were treated with antibiotics to generate germfree *Hydra*. These germfree polyps were recolonized in mono-association with one of the 5 main colonizer and either treated with phage TJ1 or with SM-Buffer as a control. The bacteria were quantified via plating and counting the CFU/polyp and the phages were quantified via qPCR. **B:** Mean CFU/polyp (± SE) over time of *Curvibacter* sp., *Duganella* sp., *Undibacterium* sp., *Acidovorax* sp. or *Pelomonas* sp. on *Hydra* with phages (Δ) or without (•) (n=5). Stars (*=p<0.05, **= p<0.01) indicate significantly difference between phage treatment and control (One-way-ANOVA). **C:** Bar plot of deltaCt values of Curvibacter (blue) and phage (grey) over time in the phage TJ1 treated polyps monocolonized with *Curvibacter* sp., *Duganella* sp., *Undibacterium* sp., *Acidovorax* sp. or *Pelomonas* sp. (n = 5). Stars (*=p<0.05, **= p<0.01) indicate significantly difference between the time of incubation in comparison to 0 h (GLM).

Similar broad host ranges (polyvalency) have been described for other phages that belong to the *Myoviridae* (Goodridge *et al.*, 2003) and other freshwater phages (Malki *et al.*, 2015). Polyvalency has been suggested to be linked to poor-nutrient conditions as an adaptation to low host cell numbers in aquatic environments (Chibani-Chennoufi *et al.*, 2004) and is also wide spread in bacteriophages that infect *Burkholderiales* (Langley *et al.*, 2003; Kawasaki *et al.*, 2009). In this context the *Burkholderia cepacia complex* (BCC) group of bacteria is intensively studied. This group of bacteria includes beneficial and pathogenic bacteria of plants but also human opportunistic pathogens, such as *Burkholderia cenocepacia* causing pulmonary infection in the background of cystic fibrosis and chronic granulomatous disease (Mahenthiralingam *et al.*, 2005). Several phages have been characterized for their ability to control *B. cenocepacia* among those *Burkholderia* phage KS10 (Goudie *et al.*, 2008), which shared 25 of 54 proteins with phage TJ1. The presence of phylogenetic similar prophages in distantly related bacteria Betaproteobacteria as well as Gammaproteobacteria (Supplementary Fig. S2) supports our findings of a brought host range.

Our data provide direct evidence that phage TJ1 can affect the abundance of symbiotic bacteria which are stably associated with their eukaryotic host. Due to its ability to infect and kill other *Curvibacter* strains from different *Hydra* species, phage TJ1 may function as a weapon against these bacteria and secure its host the dominant position on the glycocalyx of *Hydra vulgaris* (AEP).

### *Curvibacter* can use phage TJ1 to shape the *Hydra* associated microbiota

Given the above results, phage TJ1 may contribute to the regulation of the microbial composition. However, to exert its regulatory function phage TJ1 must switch from a lysogenic to a lytic life cycle even under normal environmental conditions. According to our recently published modelling approach the lytic pathway of phage TJ1 is predicted to be favored at low frequencies of *Curvibacter sp.* and high frequencies of *Duganella sp.* (Li *et al.*, 2017). For this we thought that the presence of different bacterial strains is sufficient to exert the regulatory function of phage TJ1. To test this, we monocolonized *Hydra vulgaris* (strain AEP) with *Curvibacter* sp. and exposed them separately to five bacterial strains commonly found on *Hydra vulgaris* (strain AEP): *Curvibacter* sp. (double recolonization as control), *Duganella* sp., *Undibacterium* sp., *Acidovorax* sp. and *Pelomonas* sp.. After 16 h we quantified phages, *Curvibacter* (Fig. S4) and all bacteria (Fig. S5) by qPCR (Fig. 4A). Intriguingly, we observed a significant higher phage/*Curvibacter* ratio compared to the control treatment, when *Curvibacter* mono-associated polyps were exposed to phage TJ1 sensitive bacteria including *Duganella* (Generalized linear model (GLM), F= 6.9518, p=0.03723), *Pelomonas* (GLM, F = 6.9518, p = 0.02432) and *Undibacterium* (GLM, F = 6.9518, p = 0.00299) (Fig. 4B).

**Fig 4.**
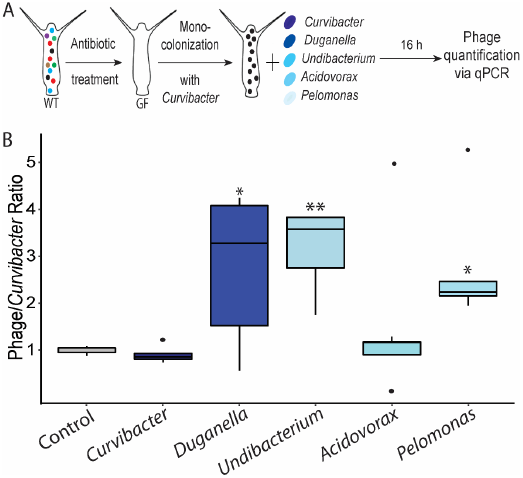
**A:** Experimental set-up; WT polyps were treated with antibiotics to generate germfree *Hydra*. These germfree polyps were recolonized in mono-association with *Curvibacter* and exposed to one of *Hydras* main colonizer (*Curvibacter, Duganella, Undibacterium, Acidovorax* or *Pelomonas*) or sterile *Hydra* culture medium as control. Phages were quantified via qPCR (n=5); **B:** Boxplot of phage/*Curvibacter* ratio after the polyps were exposed to the bacteria. Stars (*=p<0.05, **= p<0.01) indicate significantly differences between treatment and control (Generalized linear model).

The addition of bacterial strains that were resistant to phage infection, such as *Curvibacter* sp. and *Acidovorax* sp., did not lead to significant changes in the ratio between phage and *Curvibacter* sp. (Fig. 4B). An elevated prophage induction in *Curvibacter* sp. or cross-infectivity of these bacterial strains therefore can be excluded. The observation that phage/*Curvibacter* ratios increased in the presence of specific bacteria clearly demonstrates that the prophage of *Curvibacter* can exert a regulatory function on the microbiota composition.

We could show that already the addition of bacterial strains to the surrounding of monocolonized *Hydra* leads to phage replication. This might be triggered by small signal molecules released by bacteria (Ghosh *et al.*, 2009) or the ability of low abundant phages to cross-infect new colonizing bacteria. It could also be speculated that an increased abundance of bacteria stimulate the immune response of the eukaryotic host *Hydra* (Franzenburg *et al.*, 2012), which may lead to prophage induction. Nevertheless, this experiment clearly shows that phage TJ1 is active on *Hydra* without any additional artificial stimulation and interferes with microbial colonization.

### Impact of *Curvibacter* phage TJ1 on microbial community composition on *Hydra*

To validate the impact of phage TJ1 on a bacterial community level, we exposed polyps that were recolonized with *Duganella, Undibacterium, Acidovorax* and *Pelomonas* to phage TJ1. After 24 h and 72 h we analyzed the bacterial community composition by 16S rRNA amplicon sequencing and observed a reduction in the relative abundance of *Duganella* (GLM, F= 14.077, 72 h: p=0.0233) and *Pelomonas* (GLM, F= 14.077, 24 h: p= 0.0064, 72 h: p<0.001) in the phage treatments compared to the control (Fig 5A). This reduction was accompanied by a relative increase of the phage resistant bacterium *Acidovorax* (GLM, F= 14.077, 24 h: p< 0.001, 72 h: p<0.001) (Fig. 5 B). The recolonization of *Undibacterium* was not successful and therefor absent in treatment and control. However, our experiment demonstrates for a bacterial community with reduced complexity that the presence of a temperate phage has the potential to cause tremendous shifts in the bacterial community composition. Considering that all host associated bacterial communities also feature a diverse phage community e.g. the human gut (Manrique *et al.*, 2016) complex phage regulated processes of the bacterial community can be expected. Our results demonstrate that already the work with bacterial isolates should consider that single bacterial strains can be associated with prophages. Environmental and bacterial frequency depend factors that affect lysogenic or lytic decision of phages can impair experimental outcomes. Moreover, cross-infection and horizontal gene transfer are additional factors that should be taken into account when studying bacteria-bacterial as well as bacteria-host interactions.

**Fig 5.**
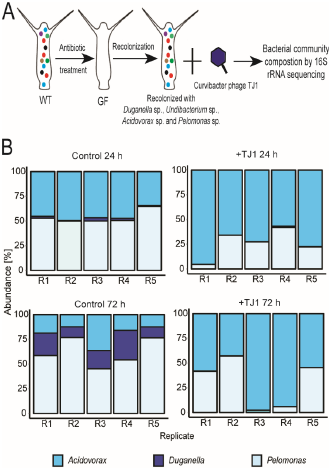
**A:** Experimental set-up; WT polyps were treated with antibiotics to generate germfree *Hydra*. Germfree polyps were recolonized with *Duganella, Undibacterium, Acidovorax* and *Pelomonas*. Recolonised polyps were exposed to Curvibacter phage TJ1. After 24 h and 72 h hours bacterial community composition was analysed by 16S rRNA amplicon sequencing (n=5). As control served recolonized polyps that were exposed to Hydra medium; **B:** Barplots illustrate the relative abundance of *Duganella sp., Undibacterium sp., Acidovorax sp.* and *Pelomonas sp.* of controls and phage treated polyps after 24 h and 72 h.

## Conclusion

This study elucidates in the early emerging metazoan *Hydra* the potential regulatory role of prophages and uncovers the capability of *Curvibacter* sp. phage TJ1 to directly affect and shape the *Hydra* associated microbiota. *Curvibacter* sp., the most dominant colonizer of *Hydra vulgaris* (AEP), is associated with a prophage which can be induced and cross-infect different bacterial strains. Observed differences in both, prophage excision from the host genome and infectivity of phage TJ1 between experiments in culture and on *Hydra* imply that the habitat of the prokaryotic host and/or bacterial frequency dependent factors influence (bacterial) host-phage interactions. In conclusion, prophages are hidden parts of the microbiome and can interfere with bacteria-bacteria interactions. Therefore prophages have the potential to influence the composition of host associated bacterial communities.

## Acknowledgements

The work was supported by grants from the Deutsche Forschungsgemeinschaft (DFG), the Collaborative Research Center 1182 (“Origin and Function of Metaorganisms” subproject A4 to SF and TB). TL gratefully appreciates support from the Volkswagen Foundation (“Experiment!- In search of bold research ideas” Az 91135). TB acknowledge support from the Canadian Institute for Advanced Research (CIFAR).

