## Supplementary material for "The neglected part of the microbiome: Prophage TJ1 regulates the bacterial community of the metaorganism *Hydra*"

**Supplementary figures**

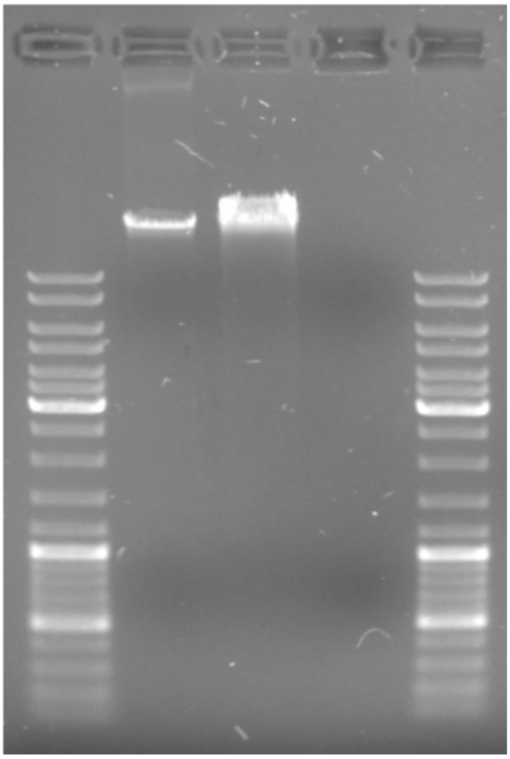

FigureS1: 0.8% Agarose gel of phage TJ1 DNASe digestion (From left to right: 1kb DNA ladder; phage TJ1 DNA; S1 nuclease digestion of phage TJ1 DNA; Turbo DNAse digestion of phage TJ1 DNA; 1kb DNAladder)

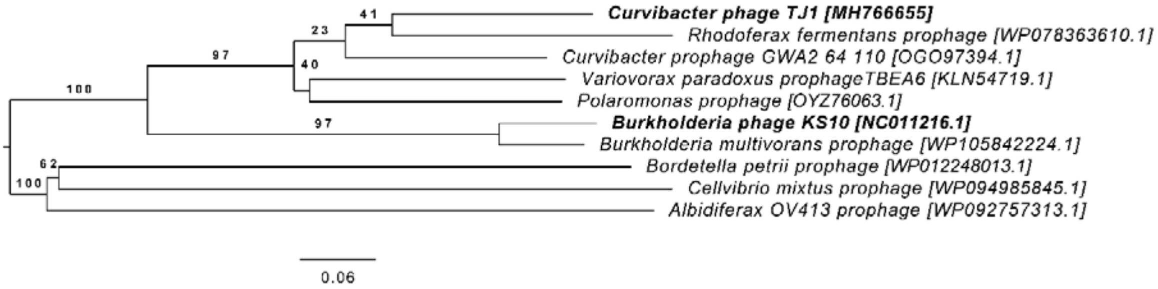


Figure S2: Phylogenetic analysis of *Curvibacter phage TJ1*, *Burkholderia phage KS10* and other prophages. Prophage sequences were identified by PHASTER (Phage Search Tool Enhanced Release) in bacterial genomes featuring proteins highly similar to Curvibacter phage TJ1. Phylogenetic tree was calculated by VICTOR using the GENOME-BLAST Distance Phylogeny method (GBDP). Branch length are scaled in terms of the GBDP distance formula $d_0$. Average support values are presented in percent.

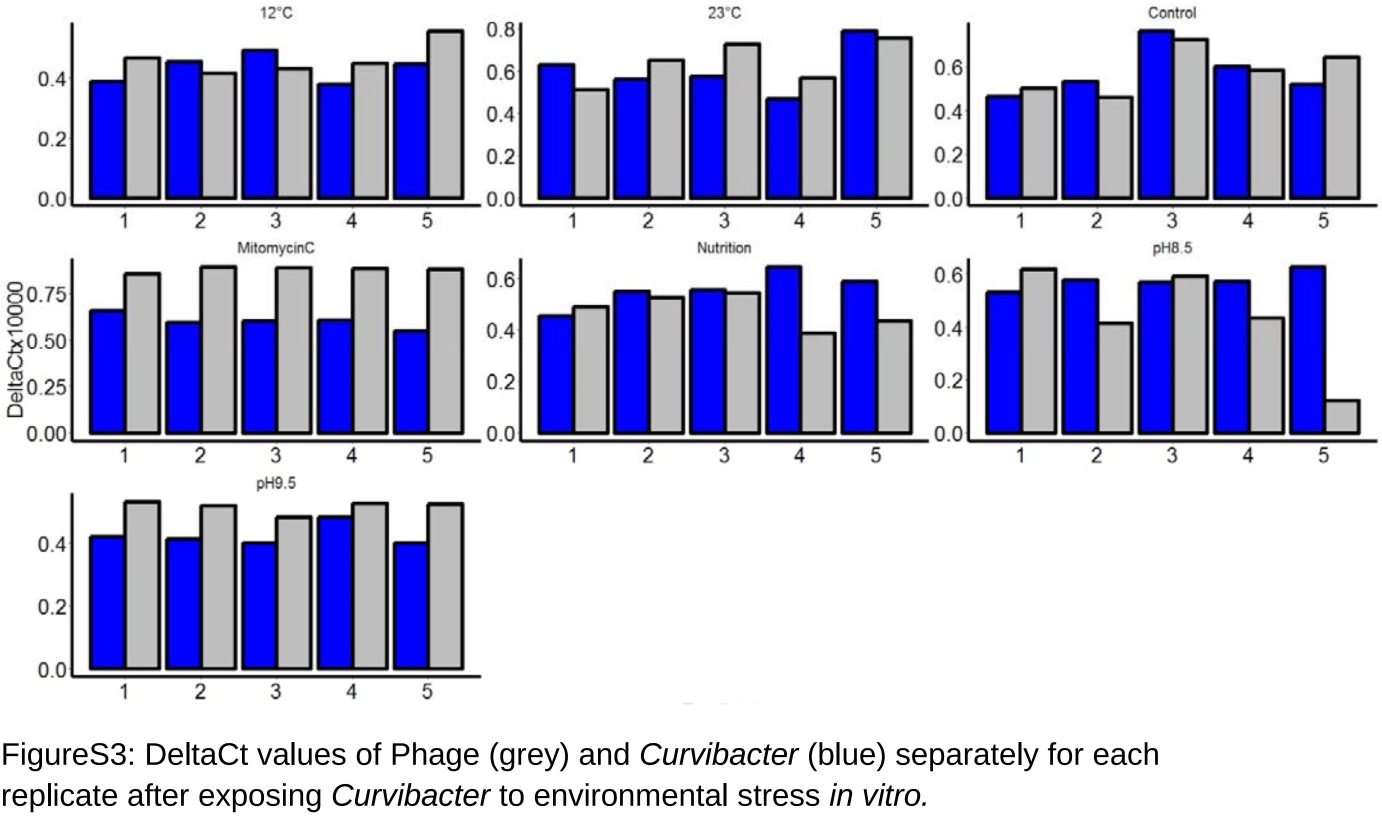


FigureS3: DeltaCt values of Phage (grey) and *Curvibacter* (blue) separately for each replicate after exposing *Curvibacter* to environmental stress *in vitro.*

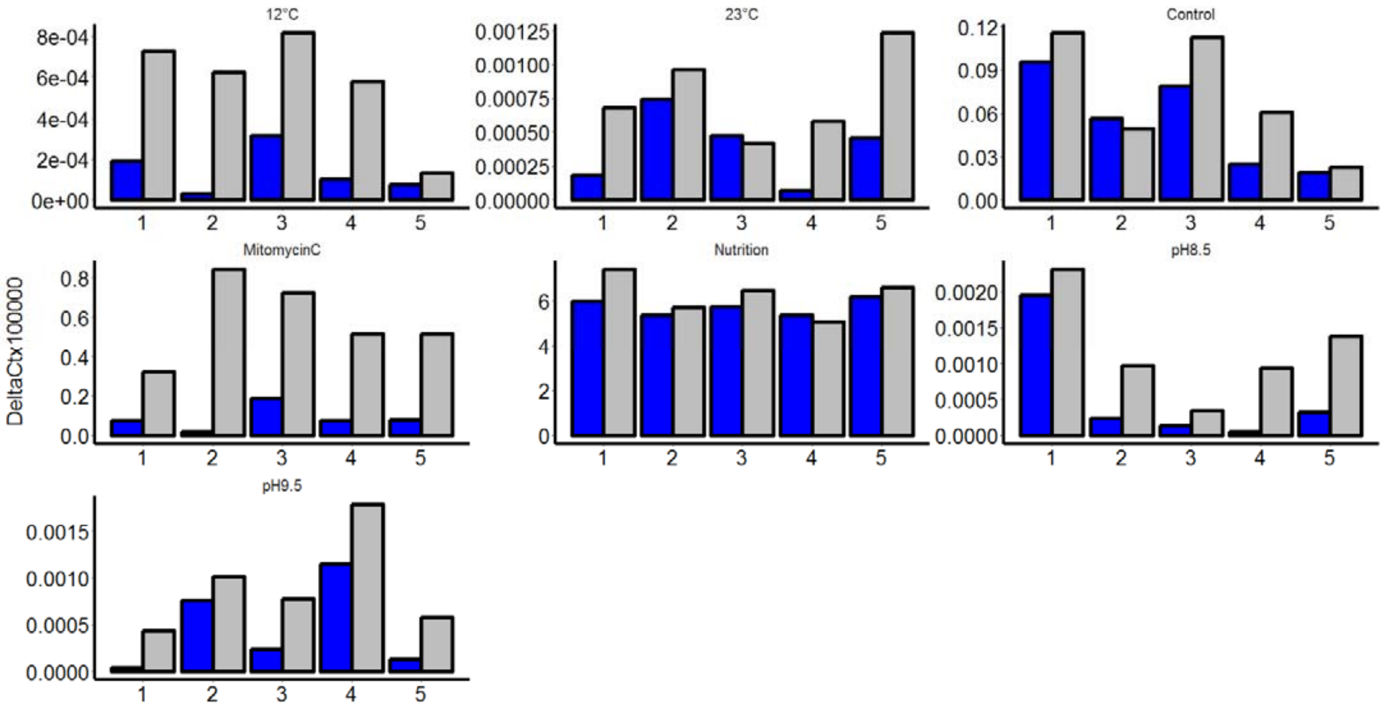


FigureS4: DeltaCt values of Phage (grey) and *Curvibacter* (blue) separately for each replicate after exposing *Curvibacter* mono-colonized *Hydra* to environmental stress *in vivo.*

Table S1: ORFs located within phage TJ1 of Curvibacter sp.

| ORF | Strand | Coding region (bp) LeftEnd | RightEnd | Protein size (aa) | Possible function | hmm name | blastp results | GenBank accession no. |
|---|---|---|---|---|---|---|---|---|
| 1 | - | 365 | 1594 | 409 | Putative capsid assembly protein F | Phage_Mu_F | *Curvibacter sp. GWA2 64 110* | OGO97394.1 |
| 2 | - | 1587 | 1910 | 107 | Uncharacterized protein | DUF4406 | *Curvibacter sp. GWA2 64 110* | OGO97393.1 |
| 3 | - | 1907 | 2059 | 50 | Uncharacterized protein | | *Vibrio phage vB VpS PG07* | AXQ66778.1 |
| 4 | - | 2066 | 2347 | 93 | Uncharacterized protein | | *Herbaspirillum sp. K1R23-30* | WP_119769014.1 |
| 5 | - | 2360 | 3919 | 519 | Portal protein | DUF935 | *Curvibacter sp. GWA2 64 110* | OGO97391.1 |
| 6 | - | 3916 | 5508 | 530 | Uncharacterized protein | | *Curvibacter sp. GWA2 64 110* | OGO97390.1 |
| 7 | - | 5508 | 6017 | 169 | Uncharacterized protein | DUF1804; HTH_23; UPF0147 | *Curvibacter sp. GWA2 64 110* | OGO97389.1 |
| 8 | - | 6021 | 6326 | 101 | Uncharacterized protein | | *Rhodoferax fermentans* | WP_078363610.1 |
| 9 | - | 6323 | 6535 | 70 | RNA polymerase-binding transcription factor DksA | zf-dskA_traR; HrpB7; ZF-HD_dimer | *Curvibacter sp. GWA2 64 110* | OGO97387.1 |
| 10 | - | 6538 | 7056 | 172 | Uncharacterized protein | | *Pandoraea apista* | WP_052765636.1 |
| 11 | - | 7053 | 7775 | 240 | Membrane-bound lytic murein transglycosylase F | SLT | *Curvibacter sp. GWA2 64 110* | OGO97385.1 |
| 12 | - | 7777 | 8121 | 114 | Uncharacterized protein | Holin_2-3; PsbJ | *Polaromonas sp.* | OYZ76063.1 |
| 13 | - | 8220 | 8642 | 140 | Uncharacterized endonuclease HI_1296 | SNase | *Hydrogenophaga sp. H7* | OPF63875.1 |

Table S2: Host range of *Curvibacter* phage TJ1 on *Hydra* associated bacteria. ✔ indicates a plaque formation and an infection of the bacterium with phage TJ1, while ✖ indicates no plaque formation and resistance to phage TJ1 infection.

| Class | Order | Family | Genus | Phage TJ1 | |
|---|---|---|---|---|---|
| β-Proteobacteria | *Burkholderiales* | *Comamonadaceae* | *Curvibacter* AEP 1.3 | ✖ | |
| β-Proteobacteria | *Burkholderiales* | *Comamonadaceae* | *Curvibacter* Mag 1.1 | ✔ | 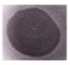 |
| β-Proteobacteria | *Burkholderiales* | *Comamonadaceae* | *Curvibacter* Hvul | ✔ | 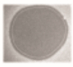 |
| β-Proteobacteria | *Burkholderiales* | *Oxalobacteraceae* | *Undibacterium* C1.1 | ✔ | 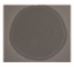 |
| β-Proteobacteria | *Burkholderiales* | *Comamonadaceae* | *Acidovorax sp.* AEP 1.4 | ✖ | |
| β-Proteobacteria | *Burkholderiales* | *Comamonadaceae* | *Pelomonas sp.* AEP2.2 | ✖ | |
| β-Proteobacteria | *Burkholderiales* | *Oxalobacteraceae* | *Duganella* C1.2 | ✔ | 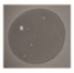 |
| β-Proteobacteria | *Burkholderiales* | *Oxalobacteraceae* | *Duganella* Oli 1.1 | ✔ | 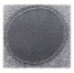 |
| ϒ-Proteobacteria | *Pseudomonaceae* | *Pseudomonas* | *Pseudomonas* AEP 1.2 | ✖ | |
| ϒ-Proteobacteria | *Pseudomonaceae* | *Pseudomonas* | *Pseudomonas* Mag 2.2 | ✖ | |
| ϒ-Proteobacteria | *Pseudomonaceae* | *Pseudomonas* | *Pseudomonas* Oli 1.2 | ✖ | |
| Bacteriodetes | *Flavobacteriaceae* | *Flavobacterium* | *Flavobacterium* Mag 1.4 | ✖ | |
| β-Proteobacteria | *Neisseriales* | *Neisseriaceae* | *Vogesella sp* AEP 1.1 | ✔ | 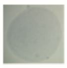 |

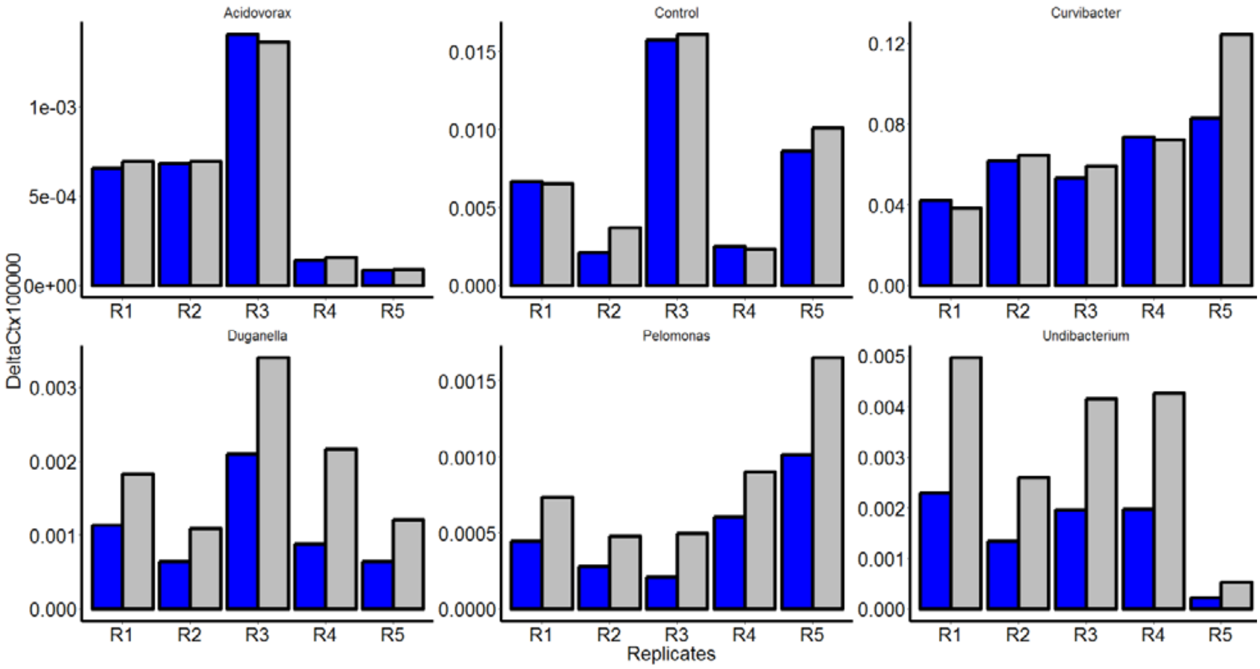


FigureS4: DeltaCt values of Phage (grey) and *Curvibacter* (blue) for the different replicates after exposing *Curvibacter* mono-colonized *Hydra* to other bacterial colonizer.

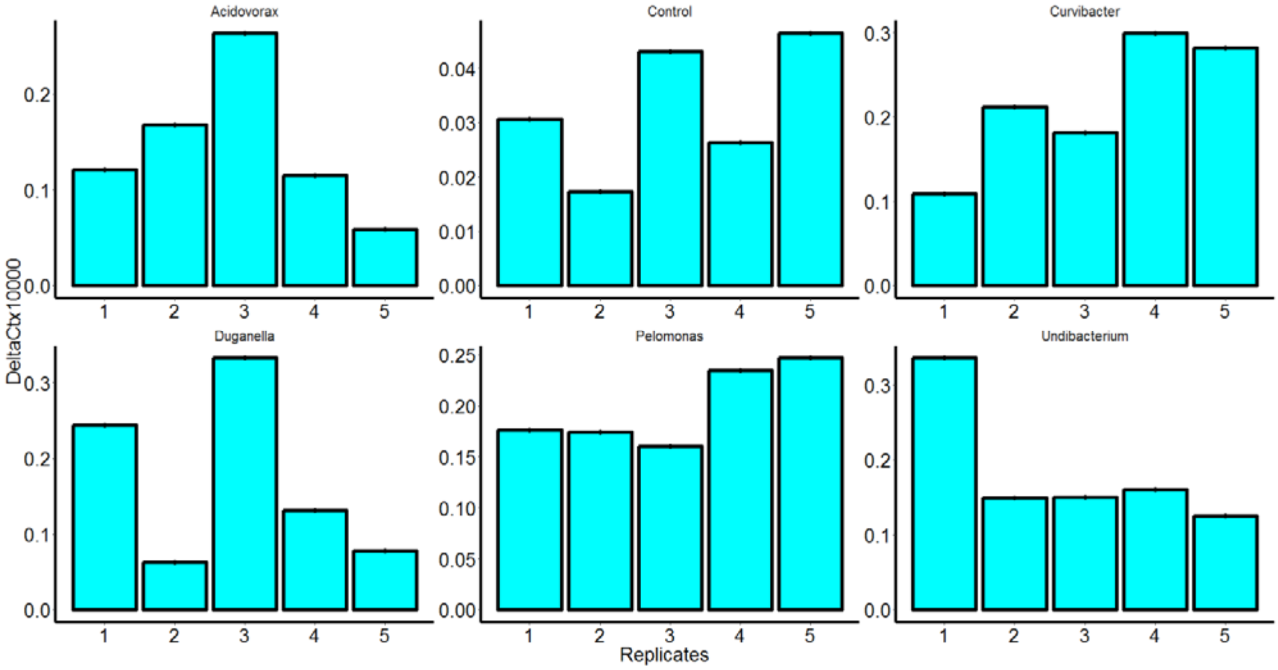


FigureS5: DeltaCt values of the Eubac Primer in the different replicates after exposing *Curvibacter* mono-colonized *Hydra* to other bacterial colonizer.
